# Biological and ecological insights from a rare underwater footage of the white shark (*Carcharodon carcharias*) in the Mediterranean Sea

**DOI:** 10.64898/2026.06.15.732360

**Authors:** Carlo Cattano, Claudia Mininni, Derk Remmers, Mario Santoro, Marco Milazzo

## Abstract

The white shark *Carcharodon carcharias* (Linnaeus, 1758) is one of the most iconic and threatened marine predators globally and is listed as critically endangered by the IUCN in the Mediterranean Sea, where populations are primarily affected by fishing pressure and the degradation of critical habitats. Available information on Mediterranean white sharks mostly relies on fishing interactions and opportunistic surface records, thereby limiting evaluations on the species ecology. Here, we report an exceptionally rare underwater encounter with this species, documented by divers involved in an expedition aimed at removing ghost fishing gear from a World War II shipwreck in the Strait of Sicily (SoS), central Mediterranean. Given the exceptional nature of this observation, we analysed the video footage in detail with the aim of providing morphological information and novel insights into the species’ ecology. The footage analysis also allowed close observations on behavioural aspects and on interspecific and parasitic associations, thus providing information that is rarely achievable through surface sightings or from captured individuals, which historically represent the majority of records for this species in the region. To contextualize this observation within the SoS—an area suggested to function as a nursery and reproductive ground for the species—we also provide an updated compilation of records reported in the scientific literature over the last decade, adding 18 new records to existing datasets. This study highlights the value of shipwrecks as observation sites for rare and elusive species, and underscores the importance of non-invasive, video-based approaches in advancing knowledge of a species whose ecology in the Mediterranean remains poorly understood.

## Introduction

The white shark *Carcharodon carcharias* (Linnaeus, 1758) is one of the most charismatic shark species worldwide, whose negative reputation has historically shaped human perception of this important apex predator. It is a highly mobile, cosmopolitan species occurring up to 1300 m depth between subpolar and subtropical regions across all major ocean basins (Ebert & Dando, 2020). Several studies conducted over the past decades have substantially advanced scientific understanding of its biology and ecology, providing direct evidence of the species’ importance in regulating ecosystems and food webs (e.g. Dedman et al., 2024; Hammerschlag et al., 2025). However, studying this species remains inherently challenging due to its low abundance and elusive behaviour.

The global population of the white shark has been estimated to have declined by 30–49% over three generations (159 years), although some signs of recovery have been reported in certain areas following the implementation of conservation measures (Rigby et al., 2022). However, the scientific community agrees on the overall low population size (Huveneers et al., 2018), due to low reproductive potential and limited capacity for rapid population recovery, which makes this species extremely vulnerable to anthropogenic pressures, particularly fisheries and habitat degradation (Rigby et al., 2022). Consequently, *C. carcharias* is classified as Vulnerable (VU) in the IUCN Red List global assessment (Rigby et al., 2022), while it is listed as Critically Endangered (CR) in the Mediterranean region (Dulvy et al., 2016). The Mediterranean white sharks constitute a genetically distinct population likely shaped by historical isolation and limited connectivity with other populations (Leone et al., 2020). The species is extremely rare in the basin and presently there is a large knowledge gap on its distribution and on potential aggregation (i.e., high occurrence) sites at regional level (Moro et al., 2020; Ferretti et al., 2024). However, available evidence indicates a marked decline of the Mediterranean population since the second half of the 20th century, which has been estimated at around 61% (Moro et al., 2020), although this estimate remains highly uncertain.

A reliable estimate of population size is hindered by the nature of the available records which mostly rely on fragmented and opportunistic observations, incidental captures, and occasional reports — providing only limited insight into species ecology, population structure, and spatial dynamics (Boldrocchi et al., 2017; De Maddalena & Heim, 2012). Current knowledge on Mediterranean white sharks relies on approximately 700 records collected over the past 150 years, allowing the identification of a few putative hotspots, including the Adriatic Sea and the Strait of Sicily (SoS) (De Maddalena & Heim, 2012; Boldrocchi et al., 2017; Moro et al., 2020). However, it should be noted that the number of reported occurrences may be biased, as species identification is often challenging due to its similarity to other lamnids, such as the shortfin mako (*Isurus oxyrhinchus*) and the porbeagle shark (*Lamna nasus*), especially during the juvenile stage and when validation relies on images (Soldo, 2026; Jambura et al., 2021).

From a habitat perspective, the SoS is characterized by a heterogeneous seascape of high biological and conservation value, recently designated as an Ecologically or Biologically Significant Area (EBSA) under the Convention on Biological Diversity (CBD-COP 12 Decisions, 2014) (e.g. Di Lorenzo et al., 2017; Consoli et al., 2021). The area hosts among the highest diversity of elasmobranch species in the Mediterranean Sea (Dulvy et al., 2016) and has been identified as strategically important for the conservation of several threatened shark species, as highlighted by the designation of multiple Important Shark and Ray Areas (ISRAs) by the IUCN Shark Specialist Group. The “Strait of Sicily and Tunisian Plateau” ISRA (IUCN SSC Shark Specialist Group, 2023) represents a recognised reproductive area for *C. carcharias*, based on the temporal occurrence of mature and pregnant females, young-of-the-year (YOY) and juvenile individuals on the continental shelf waters of Sicily, Tunisia, Lybia and Malta (Saidi et al., 2005; De Maddalena & Heim, 2012; Bradai & Saidi, 2013; Boldrocchi et al., 2017; Tiralongo et al., 2020; Jambura et al., 2021). The SoS is nonetheless one of the most intensively exploited marine regions in the Mediterranean, subject to high fishing pressure from multiple fleets and gear types (Carpentieri et al., 2021). This overlap between fishing grounds and white shark hotspots increases the likelihood of incidental captures, representing a persistent threat to the species that continues to be landed in several fishing ports (Milazzo et al., 2021).

In recent years, this raising conservation concern has driven dedicated surveys specifically targeting white sharks in the SoS. However, these efforts have failed to produce any direct observations (e.g. Micarelli et al., 2025; Ferretti et al., 2024), with the presence of the species instead inferred from environmental DNA (Janrette et al., 2023).

In this context, direct underwater observations of *C. carcharias* in the Mediterranean remain virtually absent from the literature, with available records almost exclusively based on surface sightings or interactions with fishing operations (Boldrocchi et al., 2017; Moro et al., 2020).

Here, we report a detailed analysis of a rare underwater encounter with a white shark recorded by a scuba diver in proximity to a WWII shipwreck in the Strait of Sicily, central Mediterranean Sea. While the documentation of the species in this area is scientifically relevant, this study is not intended merely to add a further occurrence record. Therefore, in addition to contextualize the sighting within recent records of *C. carcharias* in the area, we rather provide a detailed description of the observation, including morphological, ecological, and behavioural notes derived from video analysis, and discuss its potential relevance for future research on this threatened species.

## Material and Methods

The encounter with the white shark occurred on 13th May 2026 during an expedition led by the Healthy Seas Foundation, in collaboration with Ghost Diving and the Society for the Documentation of Submerged Sites (SDSS), aimed at removing ghost nets in selected WWII shipwrecks of the SoS. The expedition also felt in the frame of an ongoing scientific collaboration of the authors (MM and CC) with SDSS aimed at monitoring several submerged wreck sites in the Strait of Sicily using fixed cameras, Baited Remote Underwater Video systems (BRUVs), eDNA and acoustic receivers, assessing the importance of these habitats as potential shark aggregation areas (Cattano and Milazzo in prep.)

At 10:40 am, during the dive descent of three divers towards the target shipwreck for starting the programmed ghost net retrieval, the author DR filmed an individual of *C. carcharias* approaching the divers’ group at a depth of 40 m, a few meters above a shipwreck located at a depth of 56 m on sandy bottom. The highest point of the wreck rose approximately 14 m above the seabed.

Although available, the exact coordinates of the sighting site are intentionally not provided here. Instead, we report only a broad location, at the boundary between Tunisian and Italian waters in the southeastern sector of the SoS. This decision was made because the wreck appears to represent an aggregation area for highly diverse fish assemblages and could therefore attract or concentrate fishing activities. Such concern is supported by the presence of ghost nets on the wreck, which indicates previous fishing activity on the site.

The shark encounter was recorded by three different video cameras recording at the same time. The main observation derives from footage recorded by a Sony A7 IV camera and consisted of images with a total duration of 10 seconds (Video S1). The Camera was set to a resolution of 4K with the setting of XAVC S 4K, 25 frames per second and 10bit 4:2:2 and 140Mbit/second recording. The attached Lens was a Sony GM FE1.8/14 14mm superwideangle prime lens. The setup was used in an Easydive Leo3 WI housing and a 160mm Plexiglas domeport. The shark was also visible in the videos recorded by the other two action cameras (Water Wolf 2.0) that were attached to the divers’ equipment just before their deployment on the shipwreck for subsequent video recording for biodiversity surveys. However, image quality of these additional videos was significantly lower, and the footage was difficult to analyze due to the movement of the cameras (Fig. S1). Divers reported water visibility of 15-20 m, slight southward current during the observation, water temperatures of 19.3 °C at the surface and of 15.7 °C at 54 meters depth, and the thermocline between 19 and 25 m depth. Further information provided by the divers at the sighting site reports the presence of tuna swimming in the observation area. Moreover, they reported the presence of carcasses of fish and loggerhead sea turtles (*Caretta caretta*) trapped in the ghost net around the wreck, observed during several other dives carried out in 2025 and in the days following the white shark sighting. However, no carcasses were present in the net on the day of the shark sighting.

The main HD video was analysed using the free software VLC (www.videolan.org) to extract as much relevant biological and ecological information as possible from the imagery, including distinctive morphological features for species identification, sex determination, estimation of individual size (precaudal length, PCL in cm), and assessment of behaviour following the categories proposed by Martin (2007). Shark PCL and other main body measures were estimated through the free software Image J (Schneider et al., 2012) using one of the associated pilot fish as a scaling reference, based on the average length (FL) for the species reported for the Mediterranean Sea: 28 cm FL (Reñones et al. 1999). Obtained measurements were then compared with those reported by De Maddalena et al. (2003) for a white shark specimen caught in the Gulf of Lyon in 1953. Given the difference in body size between specimens, all measurements were expressed as percentages of precaudal length, allowing direct comparison of body proportions independent of body size.

The software was also used to estimate the number of ectoparasites visible across the shark body when zooming in some selected frames (Fig.S2). Counts were performed on the dorsal and caudal fins, as well as across three distinct body regions: (1) the head (from the mouth to the last gill slit); (2) the trunk (from the last gill slit to the pelvic fin); and (3) the precaudal tail (from the pelvic fin to the caudal fin origin) (Ebert and Dando, 2020).

In addition to gathering information from the video, we performed a literature review to update the records of the species in the Strait of Sicily from 2015 to 2026. This process implemented the database shared by Boldrocchi et al. (2017), which covered historical records until October 2015. The database was compiled by including exclusively records published in ISI scientific journals, retrieved by searching the keywords “white shark”, “*Carcharodon Carcharias”,* “Strait of Sicily” “Tunisia’”, “Libya”, "Sicily” and “Malta” in Google Scholar, and checking for relevant references in the published papers.

## Results

The recorded specimen was identified at species level by checking the presence of key morphological characteristics reported in Ebert and Dando (2020) including: 1) a sharp colour change from greyish dorsally to white ventrally; 2) small black eye; 3) large, curved pectoral fins; 4) large pelvic fins, larger than second dorsal and anal fins with broad white posterior margin and tip sharply demarcated; 5) upper and lower precaudal pit present; 6) large, erect first dorsal fin that origins usually over inner margin of pectoral fins; 7) presence of black axillary lobe marking; 8) caudal fin large and crescent-shaped with subterminal notch (See Supplementary video). The video recorded a white shark followed by 12 pilot fish (*Naucrates ductor;* Linnaeus, 1758) of similar size each other (Fig. 1 and Supplementary Video). The shark’s body size was estimated by two independent observers from video frames in which the individual presented its lateral profile to the diver, allowing for an acceptable perspective for reliable measurements. Using this approach, shark size was conservatively estimated at 335 cm PCL; this value is reported in Table S1 along with other morphometric estimates. Comparison with available morphometric measures in literature showed high correspondence between morphometric proportions reported, with a mean absolute difference of 1.99 % (±0.01 SD) PCL.

**Fig. 1:**
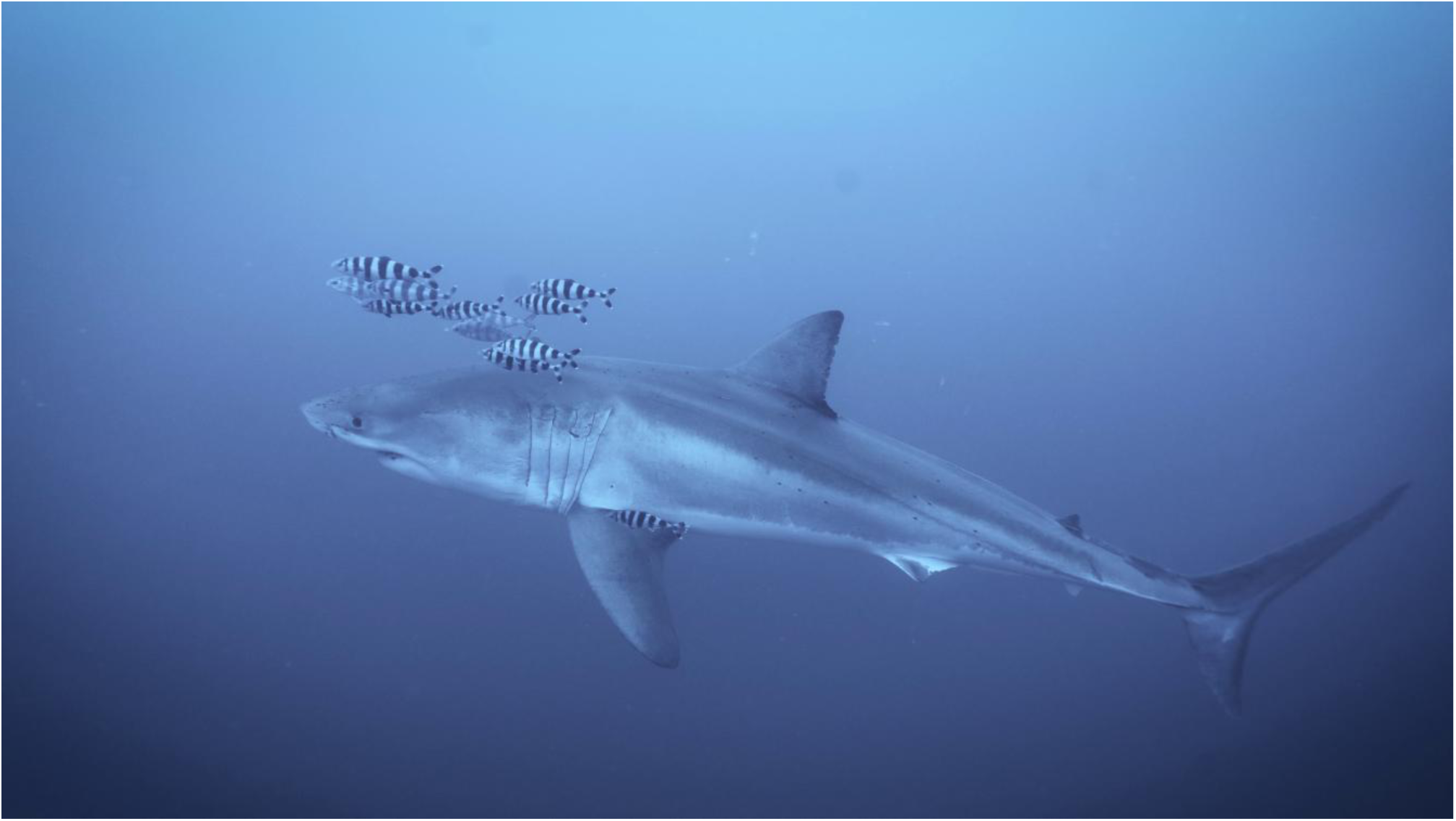
The white shark specimen recorded in the SoS the 13^th^ of May 2026 (Photo credit: Derk Remmers)

In several video frames, the claspers were visible but could not be accurately measured due to perspective distortion, nevertheless suggesting that the individual was a sexually mature male (Fig. 2). The lateral view of the individual also enabled the extraction of a still image of the dorsal fin, useful to support potential future individual identification (Fig. S3).

**Fig. 2:**
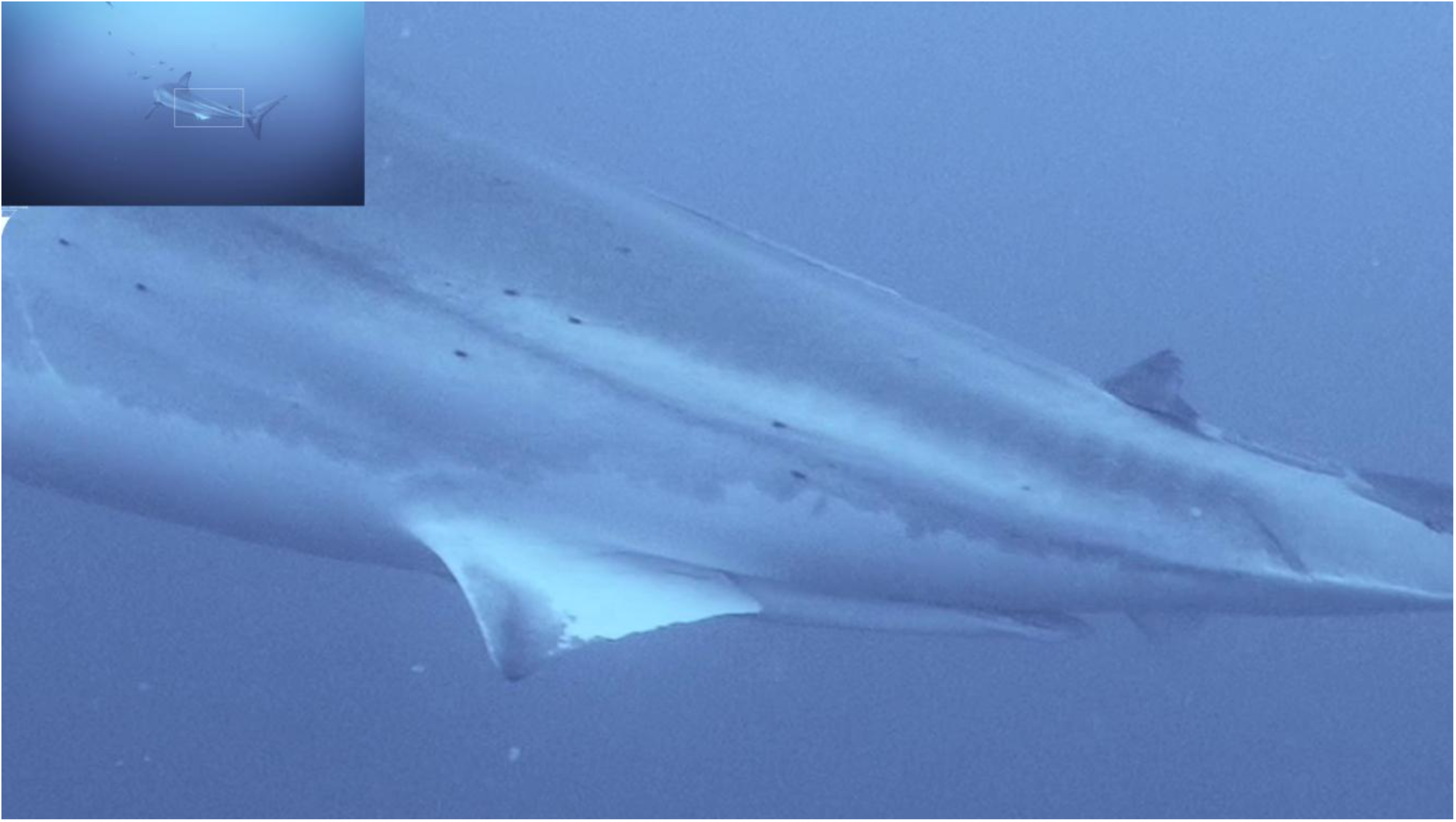
**Evidence of the presence of claspers and of copepod ectoparasites (e.g., the black dots on the shark skin) in the posterior body region of the white shark (but see Fig. S2, for further details). (Photo credit: Derk Remmers)**

What makes this encounter particularly interesting is the type of human–shark interaction that was documented, providing rare *in situ* evidence of the shark behavioural response to divers at depth. The shark initially swam towards the diver, showing pectoral fin depression and turning laterally, exposing its flank, before rapidly fleeing after the release of air bubbles by one of the divers.

Detailed examination of enlarged video frames in the laboratory revealed numerous ectoparasitic copepods distributed across the shark’s body surface, many of which displayed long posterior filaments (egg sacs), consistent with members of the family Pandaridae (Order: Siphonostomatoida Burmeister, 1835) (Fig.S2). A total of 46 parasites were counted on the lateral surface of the head, 23 on the trunk, and 9 on the posterior body region. In addition, two parasites were observed attached to the dorsal and pelvic fins.

The literature search for updated records of white shark in the Strait of Sicily yielded 18 individuals reported between 2015 and 2026 on 8 peer-reviewed publications (Supplementary Material). These records add on the 92 records reported until 2015 in Boldrocchi et al. (2017). Fifteen of these new reported individuals were from North Africa (9 in Tunisia and 6 in Libya), while the remaining three were recorded in Italian waters (2 in Lampedusa and 1 in Pantelleria islands) (Fig. 3). Sex was determined for 13 individuals (10 females and 3 males), whereas size estimates were reported for all records, ranging from 140 to 600 cm, thus spanning YOY, juveniles, sub-adults and adult individuals. Regarding record type, nine opportunistic boat-based sightings were recorded (1 in Italy, 4 in Tunisia and 4 in Libya), while the remaining individuals were captured by fishing gear: 3 by longlines, 3 by gillnets, and 3 by trawling. Interestingly, the only boat-based sighting in Tunisia regarded a live adult specimen of 500 cm (TL) trapped on the rest of a ghost fishing gear. White shark records occurred year-round, with most observations reported in spring (61%), followed by autumn (17%), and both summer and winter (11% each).

**Fig. 3.**
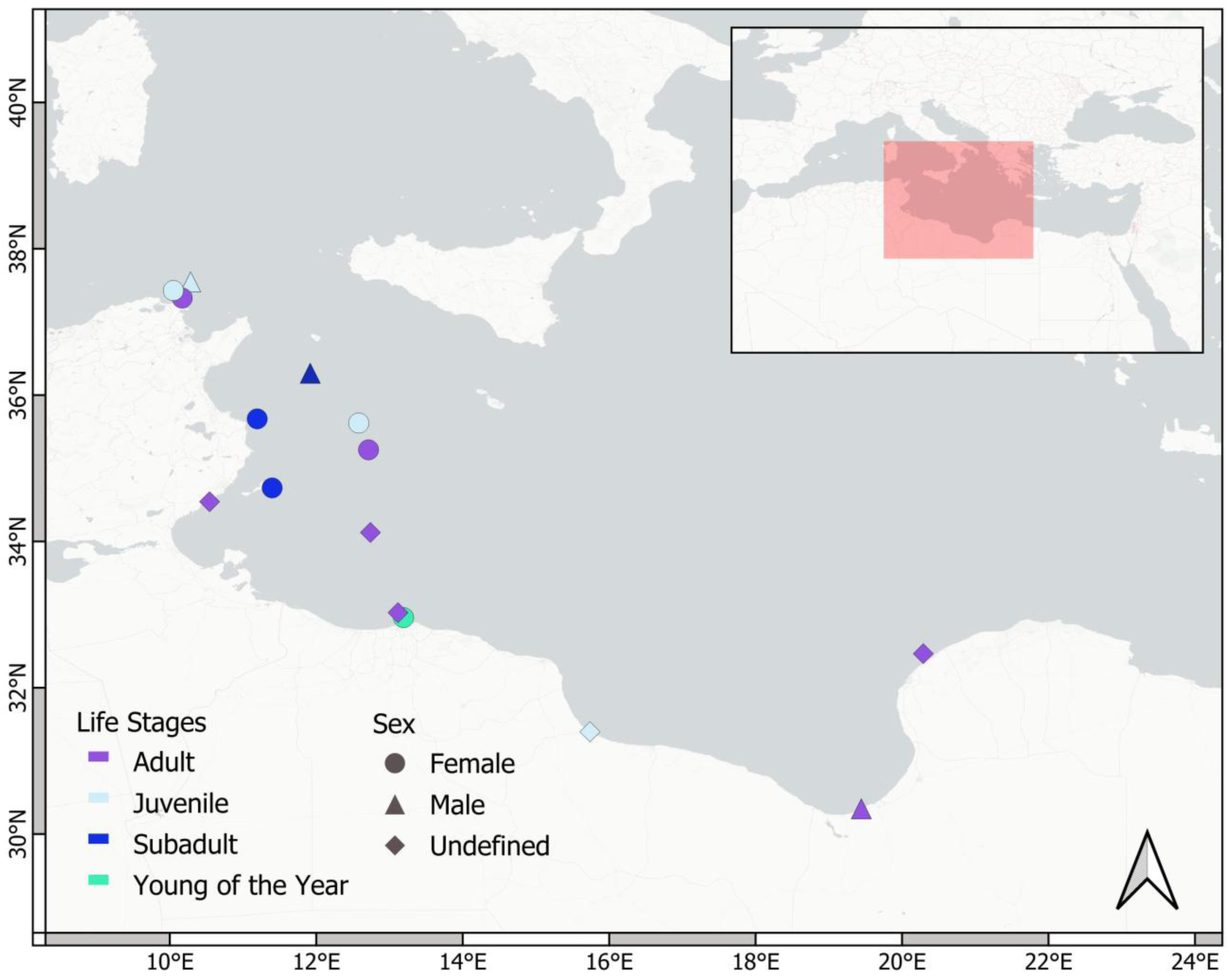
- **Distribution of the white shark records reported in scientific literature between 2015 and 2026 and categorized by sex and life stage of the observed individuals. See Supplementary Material for a complete list of the published records.**

## DISCUSSION

This study reports on the first verified underwater observation of the white shark in the Mediterranean Sea, recorded during a ghost-net removal and a scientific expedition on a WWII shipwreck of the Strait of Sicily (Central Mediterranean). Although the encounter was brief, the video footage provided valuable biological and ecological information, including reliable species identification, sex assessment, body-size estimation, behaviour during the interaction with divers, interspecific association with another fish species, and evidence of ectoparasite occurrence across the shark body.

Detailed morphometric datasets for *C. carcharias* are extremely scarce, particularly in the Mediterranean Sea, where comprehensive morphometric assessments, are generally limited to other elasmobranch species due to the rarity of whole specimens. The measurements estimated in the present study, expressed as proportions of precaudal length (PCL), showed a high degree of agreement with those reported by De Maddalena et al. (2003) for a bigger Mediterranean white shark specimen. Such low discrepancies suggest that the proportional relationships derived here provide a reliable approximation of body morphology, even when based on video observations. Minor deviations observed for some measurements are likely attributable to a combination of factors, including potential distortions associated with body orientation in the images, individual variability, and differences related to specimen condition, as the reference measurements were from a preserved museum specimen (De Maddalena et al., 2003). Differences in ontogenetic stage may also contribute to the observed slight variation This is particularly relevant for body regions that may exhibit allometric growth patterns in this species (e.g. Hunt et al., 2025), which could explain the slight variability observed in some morphometric traits among individuals of different body sizes.

The estimated size and the presence of claspers indicated that the individual reported in this study was a sexually mature adult. Males of this species have been reported to reach sexual maturity between 300 and 360 cm TL (Bruce and Bradford, 2012). The SoS has long been recognised as a key reproductive and nursery area for Mediterranean white sharks, supported by historical records of gravid females and young-of-the-year (YOY) individuals (Saidi et al., 2005; De Maddalena and Heim, 2012; Boldrocchi et al., 2017; Moro et al., 2020). In this context, the documentation of a large adult male indicates that the ecological role of the SoS is not limited to early life stages or gravid females, but may extend to multiple ontogenetic phases, reinforcing its importance as a critical habitat for the regional population (IUCN SSC Shark Specialist Group, 2023).

The white shark encounter occurred in a remote offshore area, just above a submerged shipwreck. Shipwrecks can function as artificial reefs, increasing habitat heterogeneity and complexity and supporting highly diverse benthic and pelagic assemblages (Paxton et al., 2024). These structures can concentrate high biomass of resident species and attract transient large mobile predators, including elasmobranchs, by providing localised foraging opportunities in otherwise homogeneous resource-poor offshore environments (Paxton et al., 2024). Additionally, shipwrecks may act as ecological stepping stones, enhancing ecological connectivity across seascapes. Evidence from other submersed wrecks indicates that some shark species may exhibit high site fidelity to such structures (Paxton et al., 2019). The present observation, together with additional records of other shark species at the same site during previous surveys, further supports the relevance of offshore shipwrecks in elasmobranch research and monitoring (Cattano and Milazzo, in prep.).

At a larger spatial scale, the historical occurrence of white sharks in the SoS has been linked to its distinctive biological and oceanographic features, which sustain high productivity and a complex trophic web (De Juan and Lleonart, 2010). These conditions support a high availability of white shark prey, including Atlantic bluefin tuna (*Thunnus thynnus*; e.g. Russo et al., 2021; but see Soldo and Turan 2025). Notably, divers reported the presence of large bluefin tuna in the vicinity of the study area at the time of the encounter. Although this observation remains anecdotal and lacks imagery confirmation, it is consistent with the hypothesis that local prey availability may influence the presence of adult white sharks in the area. However, the high availability of commercially important species in the area has also supported intense fishing activity, which over recent decades has increasingly targeted offshore waters of the Strait of Sicily (Paolo et al. 2024), representing an additional threat to large pelagic species, including the white shark (Ferretti et al., 2008).

Also of interest is the account provided by the divers regarding the presence, during previous and subsequent dives at the same site, of several animal carcasses, including teleost fishes and loggerhead sea turtles, entangled in the ghost net and potentially generating an odour plume.

Although this remains speculative, such cues may have contributed to increasing the likelihood of the shark approaching or investigating the wreck area. White sharks are known to rely on olfactory and other sensory cues during foraging and exploratory behaviour, and both experimental and observational studies suggest that bait, chum, and carrion can influence their attraction and behavioural responses. Notably, white sharks have been documented scavenging on large carcasses, such as whale remains, and odour stimuli associated with bait are known to affect their occurrence and behaviour during cage-diving operations (Fallows et al., 2013; Becerril-García et al., 2020).

The footage also provides relevant behavioural insights. Upon sighting the diver, the shark exhibited a moderate bilateral depression of the pectoral fins, a posture recognised as a component of agonistic displays in several shark species, including *C. carcharias* (Martin, 2007). In contrast to simple maneuvering—typically involving asymmetric and transient fin movements—sustained and bilateral pectoral fin depression is generally associated with defensive responses to perceived threats, particularly during close-range interactions, in this case with one diver of the group. However, agonistic behaviour has been described as a graded response, with the degree of pectoral fin depression apparently increasing with the perceived intensity of a threatening situation. The limited degree of fin depression observed here suggests a relatively low-intensity agonistic display within a graded behavioural response (Martin, 2007). The individual subsequently adopted a lateral orientation, exposing its flank to the diver (i.e. flank displaying; *sensu* Martin, 2007). Although not formally described as a discrete behavioural category in white sharks, such postural configuration is consistent with agonistic signaling, potentially enhancing the perceived body size of the animal during a conflict situation and often indicating an escape tendency anticipating a shift toward withdrawal. These behavioural responses may have been influenced by the calm behaviour and sustained eye contact maintained by the diver throughout the encounter, and possibly also by the diver’s apparent size and profile, which were increased by the technical equipment used for deep diving, including a rebreather, additional tanks and gear, the video camera and an underwater scooter. Indeed, the shark rapidly retreated following the release of bubbles from the open-circuit scuba regulator. Rapid withdrawal is a common component of agonistic responses and, in this case, suggests a strong sensitivity to sudden anthropogenic visual and acoustic stimuli (Martin, 2007). The absence of escalation into aggressive behaviour supports the interpretation of this interaction as defensive rather than aggressive, contributing further evidence against persistent misconceptions surrounding sharks (Mazzoldi et al., 2019). Similar response to human presence has been reported for other large sharks in the region. For example, it has been suggested that *Carcharhinus plumbeus* individuals may leave its aggregation sites under increased pressure from vessels and divers (Cattano et al., 2021), supporting the sensitivity of these species to anthropogenic disturbance.

From an ecological perspective, the video also revealed two noteworthy types of interactions: one involving a group of 12 pilot fish swimming in association with the shark near the anterior and the central portion of its body, and another characterized by the presence of numerous ectoparasites distributed across the shark’s entire body.

The association observed between the shark and the pilotfish is consistent with well-documented relationship exhibited by carangids with large pelagic predators. Such associations are generally used by the host fish for different purposes including parasite removal by chafing their bodies against the shark skin (Papastamatiou et al., 2007), concealment by other predators or prey (e.g. Cattano et al. 2025) or to enhance access to food through scavenging behavior (e.g. Ormond, 1980). An association between seven pilot fish and a white shark was recently anecdotally reported during a sighting off Libya (Jambura et al., 2021), although no detailed description was provided due to the opportunistic nature of the observation from a boat. However, documented evidence of this specie-specific interaction remains scarce in the scientific literature. The observation of pilotfish associated with a white shark reported here adds to the limited evidence of interspecific interactions involving this species, further emphasizing the ecological role of sharks in facilitating biological interactions within marine ecosystems (Cattano et al., 2025).

Another evident association involved small parasites of the family Pandaridae attached to the shark’s skin across the entire body, with their abundance decreasing from the head towards the origin of the caudal fin. The Pandaridae constitute a taxonomically complex family of ectoparasitic copepods infecting fishes, with approximately 90% of species associated with elasmobranch hosts (Cressey, 1967; Pegoraro de Macedo et al., 2023; Walter & Boxshall, 2026). The family currently includes 21 genera and 57 valid species (Walter & Boxshall, 2026). According to the recent review by Shamsi and Barton (2025), all copepod parasites reported from the skin of the white shark belong to the family Pandaridae. In the Mediterranean Sea, records of ectoparasites from white sharks are scarce and are limited to two pandarid species: *Dinemoura latifolia* from Adriatic Sea and Sicilian waters (Brian, 1906; Palomba et al., 2022), and *Echthrogaleus coleoptratus* from Portoferraio (Tyrrhenian coast of Tuscany, Italy) (Brian, 1906). Although these two species can be readily distinguished based on their morphological characteristics (Cressey, 1967), the available images do not provide sufficient resolution to allow reliable species-level identification. Furthermore, the possibility of co-infection commonly reported in other lamniform sharks (see also Santoro et al., 2025), or the occurrence of additional pandarid species cannot be excluded. This last hypothesis is further supported by the observed distribution of the parasites, which differs from previous reports. Both *D. latifolia* and *E. coleoptratus* are typically found around the pelvic, pectoral, caudal fins and caudal peduncle (Palomba et al., 2022; Shamsi & Barton, 2025), whereas in the present case most parasites were concentrated around the head region. However, these inconsistencies may also reflect the paucity of available data on the ecology of this group of parasites.

The footage analysed in this study provides valuable insights into the use of the area by the species, further supporting the importance of the SoS for white sharks. Most of the information available on *C. carcharias* in the region comes from accidental captures from industrial and artisanal fishing operations, surface observations, historical reports, or records from fish markets, often reported on social media (Boldrocchi et al., 2017; Soldo, 2026). More recently, white shark DNA has been detected at several locations in the Strait of Sicily (Janrette et al., 2023). The literature review conducted in this study, covering the last decade (2015–2026), added new records of the species to the 92 previously reported for the region by Boldrocchi et al. (2017) for the period 1937–2015. These more recent records are equally represented by fishery interactions and sightings from boats, and suggest that the area is important for all the life stages of the species. Notably, YOY and juveniles have been very recently recorded in the Libyan coasts, representing the southernmost occurrence of immature white sharks in the Mediterranean Sea, as reported by the authors (Jambura et al., 2021). The literature search also revealed that records resulted from interactions with multiple fishing gears operating in both pelagic and benthic environments, including bottom trawls, pelagic longlines, and gillnets. This evidence suggests that, although the species is primarily pelagic, it is exposed to a wide range of fishing activities, which increase its vulnerability, particularly in coastal and shelf areas. In addition, a very recent study reported a record from shallow waters in the Gulf of Gabès (Tunisia) of a live white shark entangled in lost fishing gear, exhibiting severe skin damage, with a rope cutting deeply into the muscle (Ziani et al., 2026). This record highlights the potential additional threat posed by ghost nets even to large sharks, adding to the limited evidence of interactions between ghost fishing gear and sharks in the Mediterranean Sea (Perroca et al., 2024). Notably, most of the new records occurred in spring, coinciding with Atlantic bluefin tuna migration pattern in the region (Russo et al., 2021), further supporting the association between white shark presence and this key prey species (e.g. Soldo & Turan, 2025). However, this pattern should be interpreted with caution, as it may be influenced by variation in observation effort, which has been suggested to affect the spatio-temporal distribution of records in the region (Moro et al., 2020). This work probably gained some more information than previous records for the region, providing biological information and additional ecological context that is usually missing from other Mediterranean records. Here we demonstrated how even a single, opportunistic observation can provide valuable ecological and behavioural insights that are largely inaccessible through fishery-dependent sampling. Non-invasive visual approaches, such as underwater video and BRUV systems, allow the direct observation of individuals in their natural environment, capturing behavioural and ecological information that cannot be inferred from landed specimens or incidental captures. Such approaches are therefore essential for advancing knowledge on rare and elusive species such as *C. carcharias*, even in contexts where encounters remain extremely challenging despite intense research efforts.

Understanding how this and other elusive large predators use specific habitats, such as submerged shipwrecks, is of fundamental importance. More broadly, improving knowledge of their spatial and temporal distribution through targeted field studies remains a key challenge for the effective protection of Mediterranean white sharks and of other large pelagic species of conservation concern.

## Acknowledgements

The authors acknowledge the Healthy Seas Foundation for organising and leading field activities connected to ghost-net removal at selected Mediterranean shipwreck sites, in collaboration with SDSS (Society for Documentation of Submerged Sites) and Ghost Diving. Healthy Seas is a foundation dedicated to removing ghost fishing gear and other marine litter to protect marine ecosystems, while promoting prevention, education, circular solutions and scientific collaboration. We would like to express our special thanks to Mario Arena, president of SDSS (Society for the Documentation of Submerged Sites), and Rocco Canella from the Diving Pelagos 2.0 in Lampedusa Island. We also thank Roberto Rinaldi documentarist, Dr. Federico Quattrocchi of the University of Palermo and Dr. Davide Spatafora of the Stazione Zoologica Anton Dohrn for their helpful suggestions.

---

**Table S1.** – Morphometric measures for the white shark recorded in this study and comparison with the measures reported in Maddalena et al. 2003 for a white shark whose precaudal length was 458 cm. Terminology and abbreviations from Compagno et al. 1984. N.A.: measures not available in De Maddalena et al. 2003.

| **Morphometric unit** | **Abbrev.** | **Estimated size cm (present study)** | **% PCL (present study)** | **Estimated size cm (De Maddalena et al. 2003)** | **% PCL De Maddalena et al. 2003** |
| --- | --- | --- | --- | --- | --- |
| Precaudal length | PCR | 335 | 100% | 458 | 100% |
| Pre-pectoral length | PP1 | 100 | 29.9% | 145 | 31.7% |
| Pre-first dorsal length | PD1 | 150 | 44.8% | 220 | 48% |
| Pre-pectoral pelvic space | PPS | 117 | 34.9% | 155 | 33.9% |
| Pelvic-anal space | PAS | 39 | 11.6% | 50 | 8.58% |
| Anal-caudal space | ACS | 39 | 11.6% | 50 | 10.9% |
| Body height at pelvic origin | N.A. | 46 | 13.7% | N.A. | N.A. |
| Eye length | EYL | 4 | 1.19% | N.A. | N.A. |
| First dorsal height | N.A. | 37 | 11.0% | N.A. | N.A. |
| First dorsal base | N.A. | 36 | 10.7% | N.A. | N.A. |
| Pre-orbital length | POB | 20 | 6.0% | 32 | 7.0% |
| First to fifth gill slits | HDL-PGI | 31 | 9.25% | 30 | 6.5% |
| Prebranchial length | PG1 | 85 | 25.4% | 122 | 26.6% |
| Pelvic-caudal space | PCA | 86 | 25.6% | 100 | 21.8% |
| Dorsal caudal space | DCS | 45 | 13.4% | 55 | 12% |

**Fig. S1.**
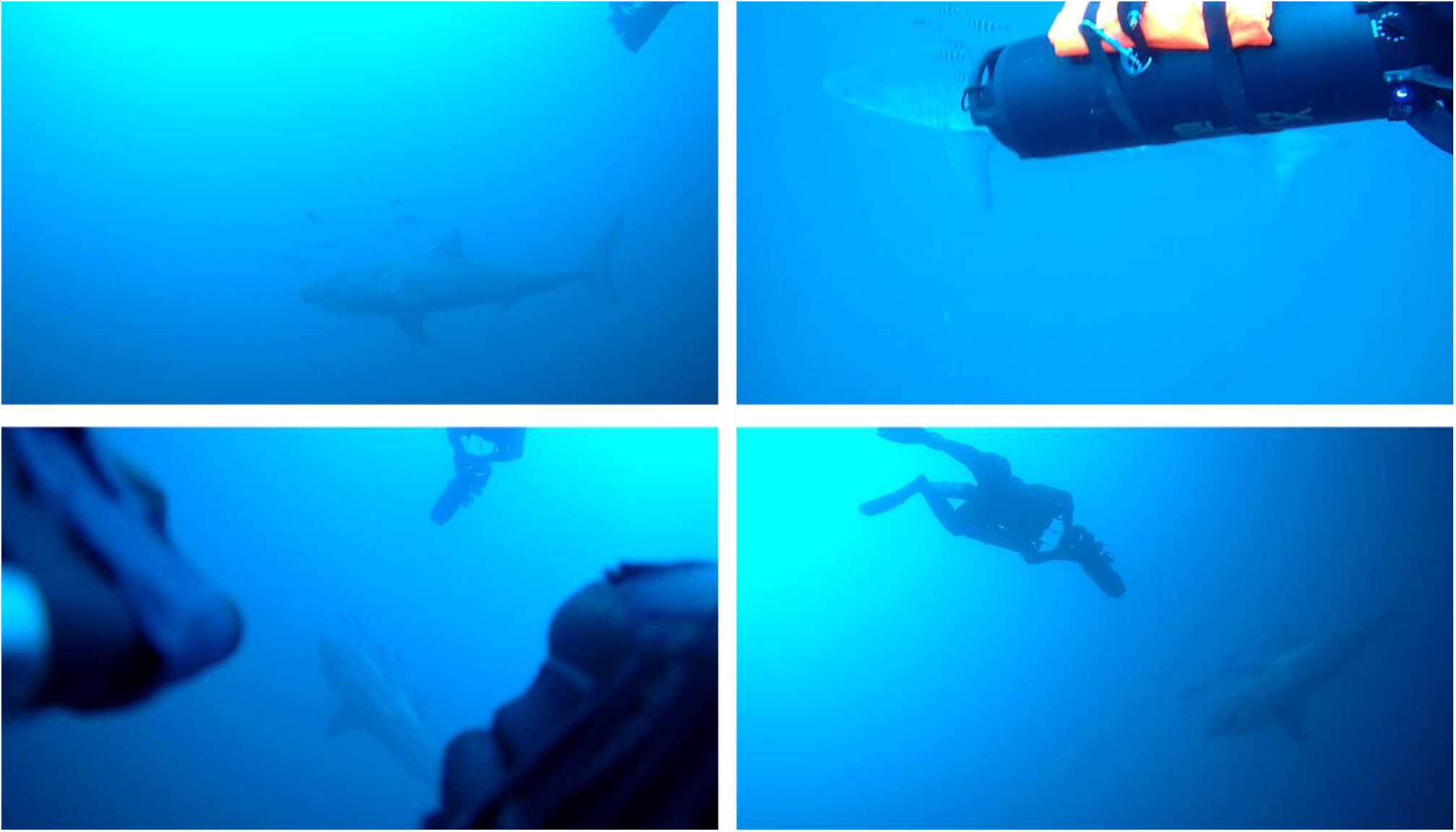
Some frames of the encounter with the white shark recorded by the other two video cameras attached to the divers’ equipment.

**Fig. S2.**
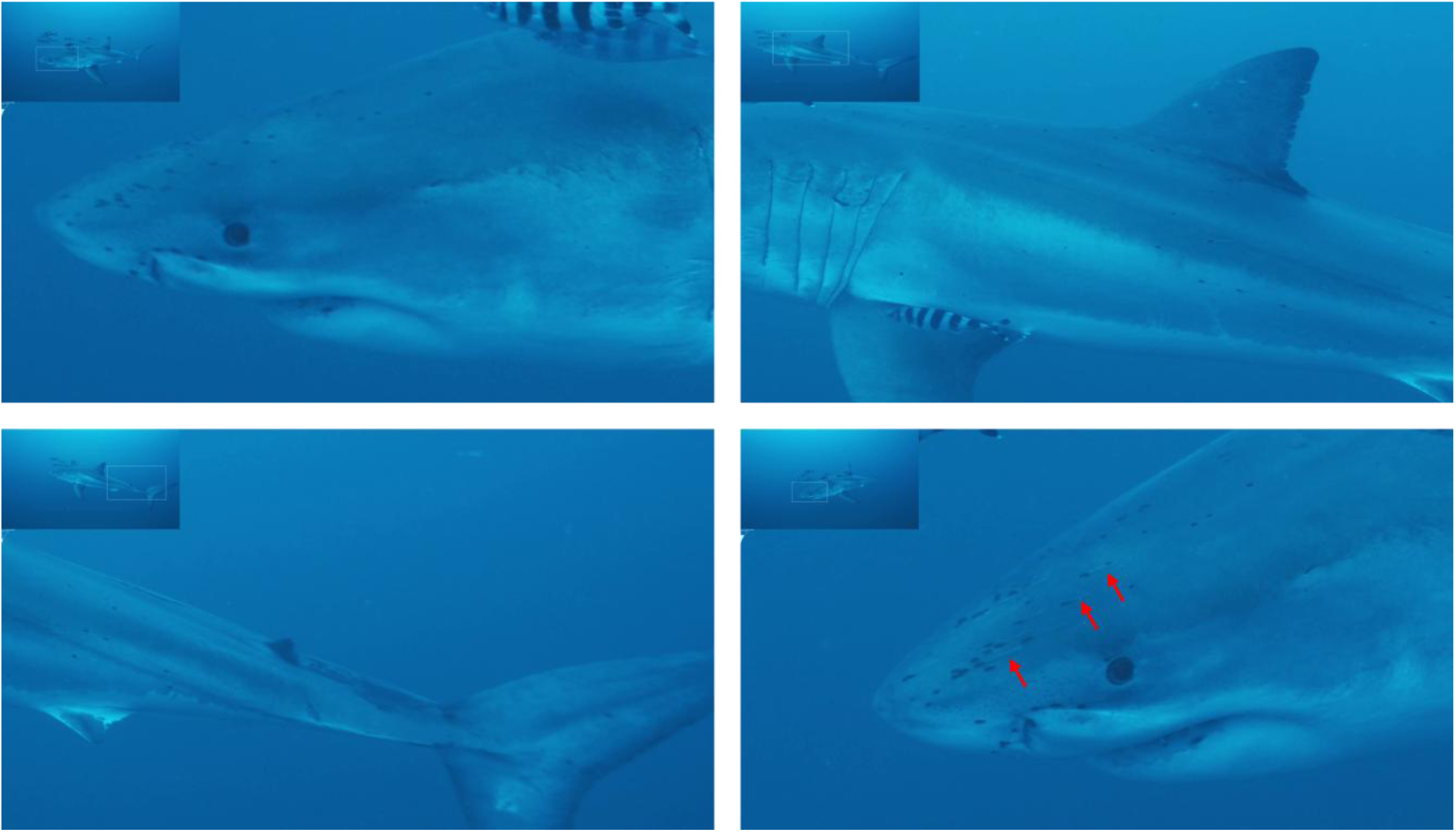
– Still frames showing the pandarid ectoparasites in the different shark body sections. Red arrows indicate some of the visible parasite egg sacs.

**Fig. S3.**
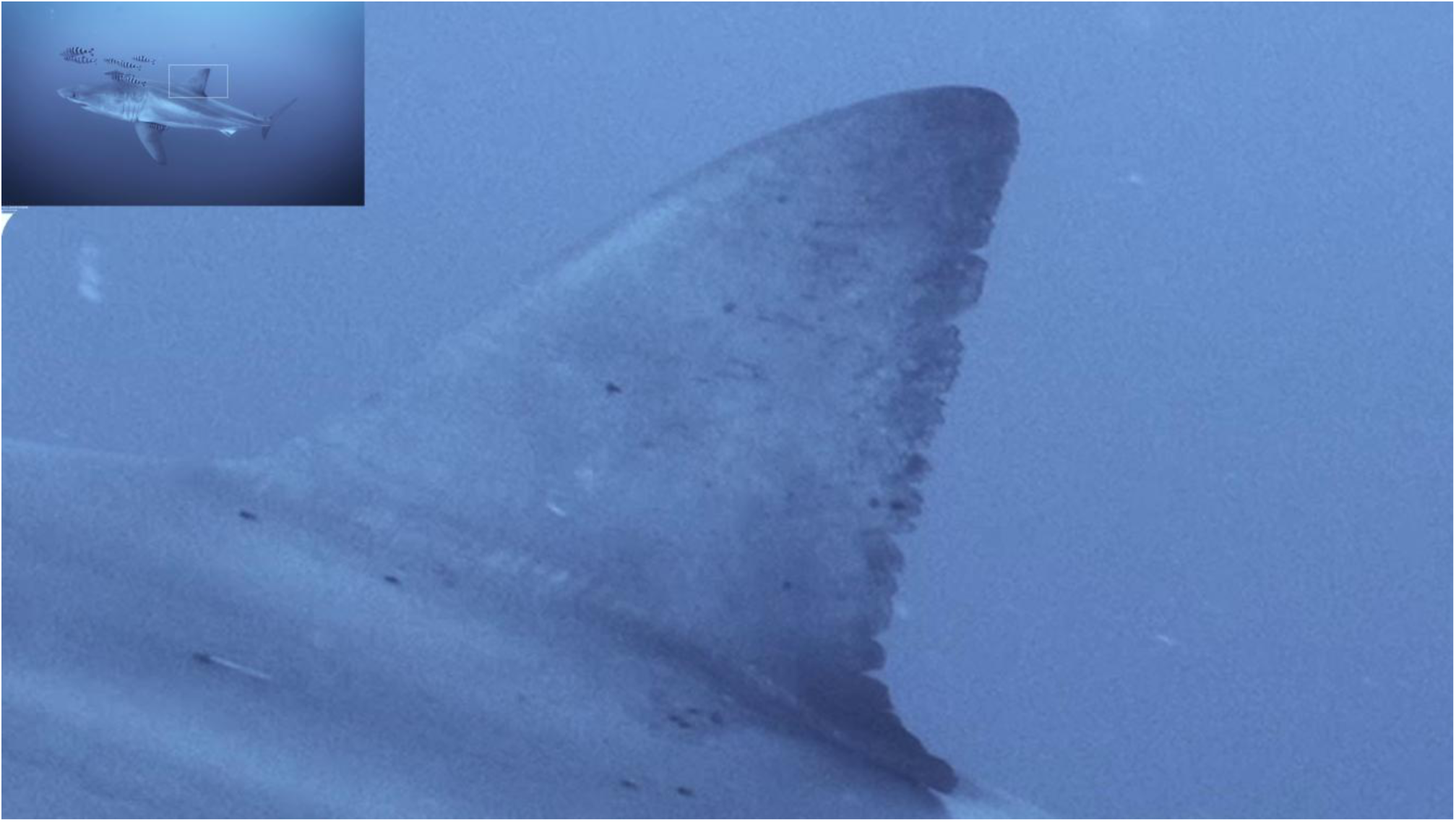
– Detail of the white shark dorsal fin. Pandarid copepods can be seen on the skin

